# A subtype of ultrasonic vocalizations during highly palatable food consumption in rats identified by machine learning–assisted classification

**DOI:** 10.1101/2025.09.09.674592

**Authors:** Koshi Murata, Yuki Ikedo, Takashi Ryoke, Kazuki Shiotani, Hiroyuki Manabe, Kazuki Kuroda, Hitoshi Yoshimura, Yugo Fukazawa

## Abstract

Identifying behavioral and physiological responses to rewarding stimuli is essential for understanding positive emotional states in animals and for investigating their neural basis. Ultrasonic vocalizations (USVs) in rats are modulated by various behavioral contexts and are thought to represent affective-like states. However, their diversity during palatable food consumption remains underexplored. In this study, we investigated acoustic features of USVs in male rats consuming chocolate, a highly palatable food. Using a machine-learning–assisted logistic regression model trained on spectrogram features, we identified a distinct USV subtype—the 40-kHz inverted-U type—that was selectively emitted during chocolate consumption. The emission of this subtype was tightly time-locked to chocolate feeding behavior. Systemic administration of naloxone, an opioid receptor antagonist, significantly reduced the emission of the 40-kHz inverted-U USVs during chocolate intake. These findings suggest that these USVs may reflect internal states associated with palatable feeding and are under modulation by the endogenous opioid system. Moreover, our data demonstrate the utility of machine learning for high-throughput, objective classification of USV subtypes. This framework provides a promising approach for decoding emotion-related vocal expressions in rats and highlights the potential of specific USVs as behavioral readouts for studying the neurobiology of positive emotions.

**Significance Statement:** Ultrasonic vocalizations (USVs) are high-frequency calls produced by rats known to correlated with emotional and motivational states, yet their diversity and functional relevance remain underexplored. In this study, we identified a novel 40-kHz inverted-U USV subtype associated with chocolate consumption. These calls were closely tied to feeding behavior and were selectively suppressed by an opioid receptor antagonist, suggesting modulation by the brain’s reward system. Using machine learning–assisted classification, we established a robust method to detect these vocalizations. Our findings highlight the potential of specific USVs as non-invasive, real-time behavioral indicators of internal states, offering new tools for studying affect, reward, and communication in animal models.

## Introduction

To understand the neurobiological mechanisms that underlie positive emotions, it is essential to identify behavioral and physiological responses to rewarding external stimuli, which in turn enables objective and quantitative assessment of such emotional states. Food consumption is accompanied by positive emotions in both animals and humans, particularly when the food is highly palatable, such as sweet-tasting foods indicating high nutritional value. These positive emotions play a key role in learning which food types to seek and in motivating animals to pursue them (Nguyen et al., 2021).

In animal studies, food palatability and associated positive emotions are frequently evaluated using preference tests, such as food choice paradigms comparing intake between two or more options (Rainwater and Güler, 2022; Cawthon and Spector, 2023). However, preference tests are indirect measures: they assess palatability and emotional valence only through differences in cumulative food intake over time. To better understand the affective component of palatable food consumption, it is important to capture the immediate behavioral and physiological responses that occur at the moment of feeding. A commonly used method for real-time assessment of positive emotions is the taste reactivity test, in which animals receive passive intraoral infusions of sweet solutions (e.g., sucrose) via implanted oral cannulas, and their facial and bodily responses are analyzed (Berridge, 2000). In this test, specific ingestive responses are defined as hedonic “liking”. However, this method is invasive and requires physical restraint for cannula connection. Moreover, it is limited to liquid taste stimuli and cannot assess emotional responses to solid foods. Therefore, more versatile, non-invasive, real-time measures of positive emotional states during food consumption in animals are needed to further investigate the underlying neural mechanisms.

Ultrasonic vocalizations (USVs) are thought to reflect positive and negative affective-like states in rats, as they occur in various behavioral contexts including social interaction, mating, play, and exposure to rewarding stimuli (Knutson et al., 2002; Portfors, 2007; Burgdorf et al., 2008). These vocalizations vary in frequency, duration, and contour shape, and may reflect the animal’s internal state or behavioral intent. Among these, 50-kHz short USVs (frequency range: 30–90 kHz, duration: 20-100 ms) have been frequently observed during approach behaviors and active engagement with stimuli, including food anticipation and consumption, suggesting an association with motivational and appetitive states (Wright et al., 2010; Wöhr, 2017). Consumption of a standard laboratory diet elicits a particular subtype of USVs; 40-kHz flat-shaped USVs (Takahashi et al., 2010; Champeil-Potokar et al., 2023). Additionally, intake of a more palatable substance (sweetened condensed milk) induces a greater number of 50-kHz short USVs during both consumption and cue-induced anticipation compared to tap water intake (Brenes and Schwarting, 2014), raising the possibility that food palatability modulates vocal output. However, it remains unclear whether specific USV subtypes with distinct acoustic features are selectively emitted during the consumption of highly palatable foods.

To address this question, we examined whether rats emit distinguishable subtypes of USVs when consuming a highly palatable food, as compared to baseline conditions without access to that food. We used sweet milk chocolate as a highly palatable stimulus, as rats exhibit a strong preference for it over standard laboratory chow, even in the absence of food deprivation (Castro and Berridge, 2014). We compared the spectrograms of USVs emitted during chocolate-access periods with those emitted during baseline periods. Using machine learning-assisted classification of spectrographic features, we constructed a logistic regression model to perform binary classification of USVs associated with chocolate presentation versus non-chocolate conditions. Through a combination of machine learning classification and conventional visual assessment, we identified a subtype of 40-kHz inverted-U USVs that were selectively emitted during chocolate consumption. Furthermore, systemic administration of the opioid receptor antagonist naloxone reduced the emission of these USVs, suggesting involvement of the endogenous opioid system. These findings suggest that food palatability may influence vocal expression in rats and raise the possibility that specific USV patterns could serve as real-time behavioral markers of emotional states during palatable feeding.

## Materials and Methods

### Ethical statements

All animal experiments were approved by the Experimental Animal Research Committee of the University of Fukui (approval numbers: R03015, R04014, R05011, and R06007). This study was performed in accordance with relevant guidelines and regulations.

### Animals

Eight-week-old male Sprague–Dawley rats were obtained from Japan SLC, Inc. and housed in our animal facility for at least 1 week before the beginning of the USV recording. A total of 64 rats were used in the experiments: 20 rats (10 pairs) for the training dataset, 22 rats (11 pairs) for the test dataset (empty group), and 22 rats (11 pairs) for the test dataset (chocolate group) followed by pharmacological experiments. The rats were housed in pairs in plastic cages (33 × 38 × 18 cm, CL-0106-2, CLEA Japan Inc.) with paper bedding (Japan SLC, Inc.) and wire cage tops. They had *ad libitum* access to pellet food (DC-8, Oriental Yeast Co., LTD.), except during USV recording sessions, with water available at all times. The rats were kept on a reverse 12–12-h light-dark cycle, with experiments conducted during the dark phase, 6 to 9 h after lights-off.

### Experimental design

First, USVs were recorded from 10 pairs of rats to create a logistic regression model which assisted visual assessment of chocolate feeding-related USV types (training dataset; Figs. 1–4). Next, an additional set of USV recordings was collected to create a test dataset (Figs. 5–8). In this phase, 11 pairs of rats were assigned to the chocolate presentation condition (chocolate group) and subsequent pharmacological experiments (Fig. 9), while another 11 pairs served as a control group that was not exposed to chocolate (empty group). The USVs from the test dataset were classified using the logistic regression model trained on the initial datasets.

**Figure 1.**
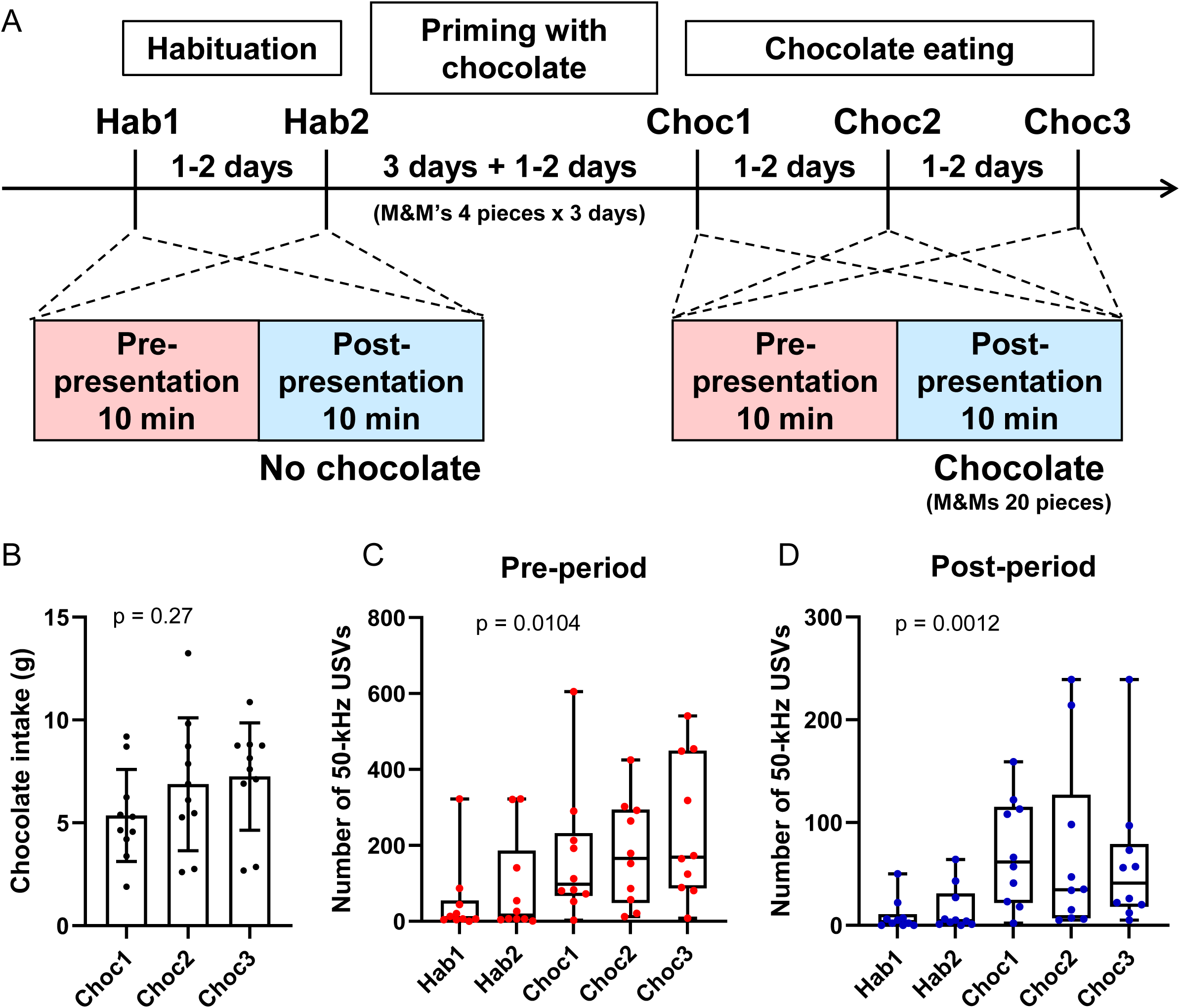
Emission of 50-kHz short USVs by pair-housed adult male rats during chocolate consumption. A: Schedule of USV recordings to obtain the training dataset for machine learning-based USV classification. Habituation sessions (hab1 and hab2) were conducted to acclimate the rats to the recording environment and to record USVs in the environment before priming with chocolate. Chocolate presentation sessions (choc1, choc2, and choc3) were conducted to record USVs before and during chocolate consumption. Each session consisted of two 10-minute periods: Pre-chocolate presentation period (pre-period) and post-chocolate presentation period (post-period). Intervals between recording sessions lasted 1–2 days. The priming phase for chocolate consumption was set between the habituation and chocolate presentation sessions, during which rats were provided with four pieces (approximately 4 g) of M&M’s milk chocolate once per day for 3 consecutive days in their home environment. The interval between the last day of priming and the first chocolate presentation session (choc1) was also 1–2 days. B: Amount of chocolate intake during the post-period. p = 0.27, one-way ANOVA. C, D: Number of 50-kHz short USVs during the pre-(C) and post-chocolate (D) periods. p = 0.0104 (C) and 0.0012 (D), Kruskal–Wallis test.

**Figure 2.**
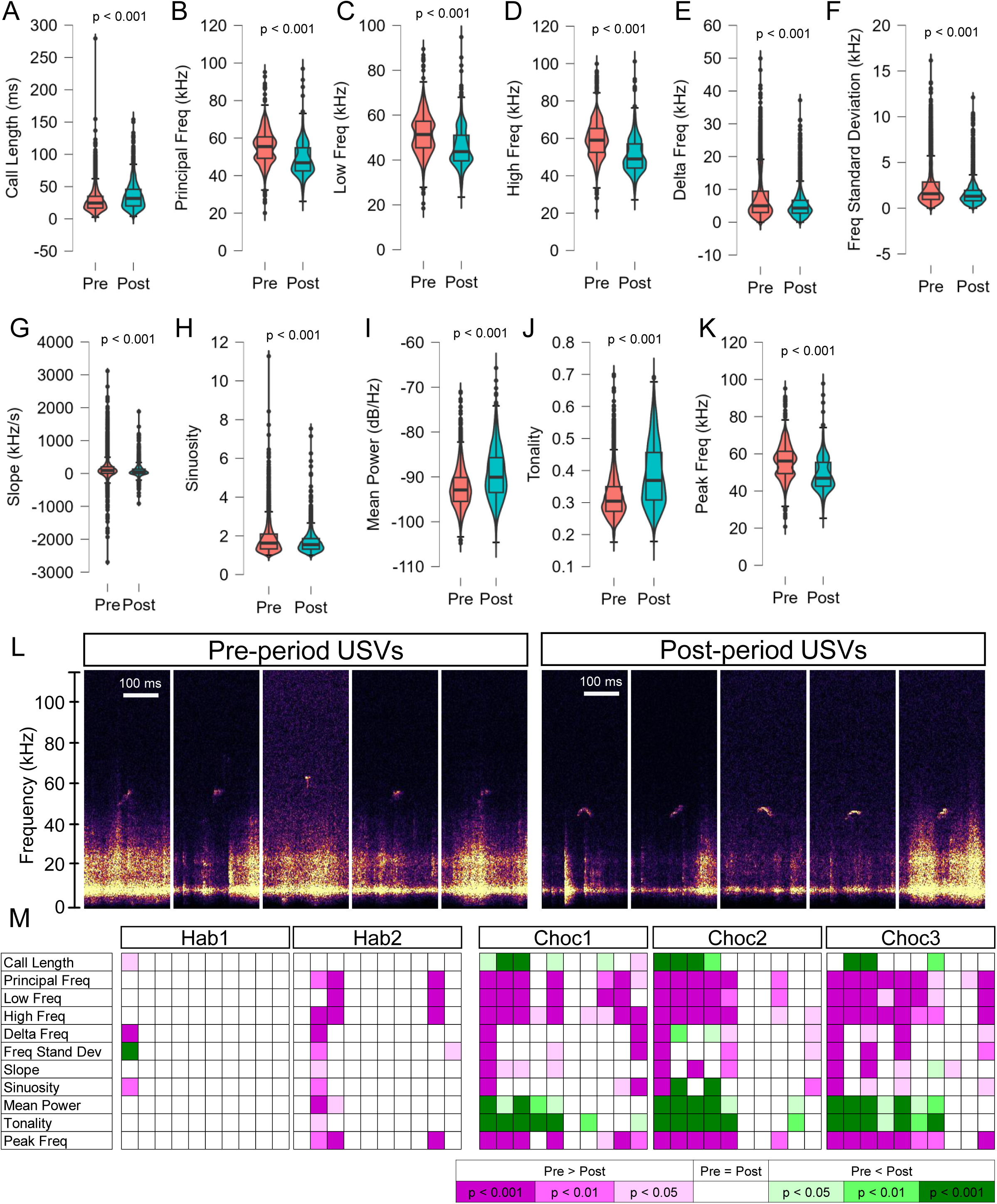
Comparison of USV spectrograms between pre- and post-periods of chocolate delivery. A-K: Violin plots representing the differences in 11 spectrogram parameters between USVs in the pre- and post-periods. USVs from a total of 10 rat pairs and three sessions (choc1, choc2, and choc3) were pooled. The total number of calls was 5,893 for the pre-period and 2,016 for the post-period. (A) Call length, (B) Principal frequency, (C) Low frequency, (D) High frequency, (E) Delta frequency, (F) Standard deviation of frequency, (G) Slope, (H) Sinuosity, (I) Mean power, (J) Tonality, (K) Peak frequency. p values were obtained from Mann–Whitney U tests. L: Representative spectrograms of the USVs from the pre- and post-periods. The five USVs with the closest Euclidean distances to median values of the 11 spectrogram parameters were selected from all USVs. M: Statistical differences in the 11 USV spectrogram parameters between the pre- and post-period are individually shown in a heatmap across rat pairs and sessions. Each cell represents the difference in a specific parameter for a particular pair. Rat pairs are sorted by the total amount of chocolate intake during the three sessions (choc1, choc2, and choc3) from largest to smallest (left to right). p values were obtained from Mann–Whitney U tests. The direction of the difference and the level of statistical significance are indicated by the color and its hue, respectively.

**Figure 3.**
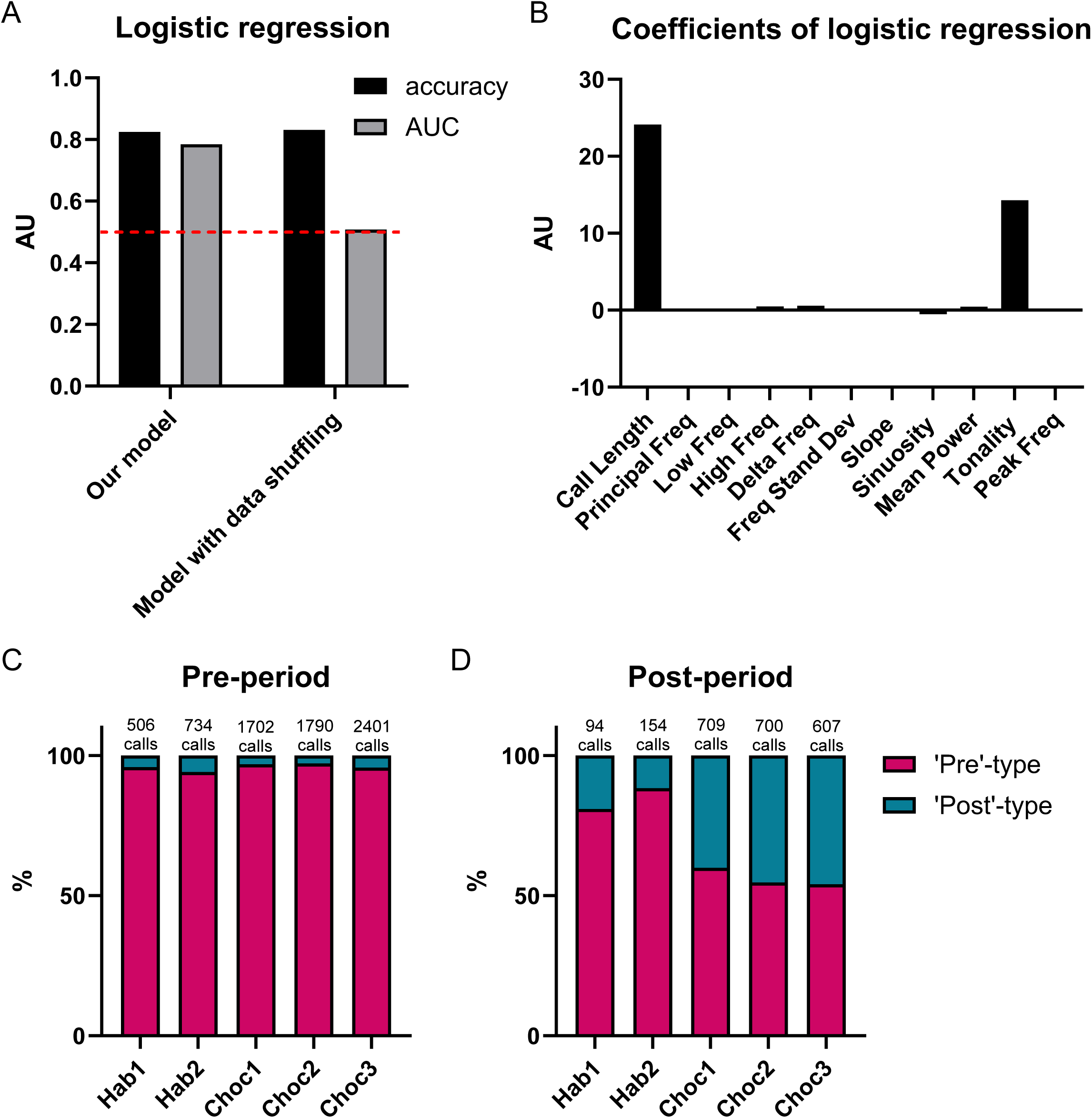
A logistic regression model for the classification of USVs into ‘pre’- and ‘post’-chocolate types. A: Evaluation of the logistic model classifying USVs into ‘pre’- and ‘post’-types, comparing accuracy and AUC with a model trained on randomized data. B: Estimated coefficients of the logistic regression model. C, D: Percentages of ‘pre’- and ‘post’-types among the total number of USVs from 10 rat pairs in the pre-(C) and post-(D) periods of each session.

**Figure 4.**
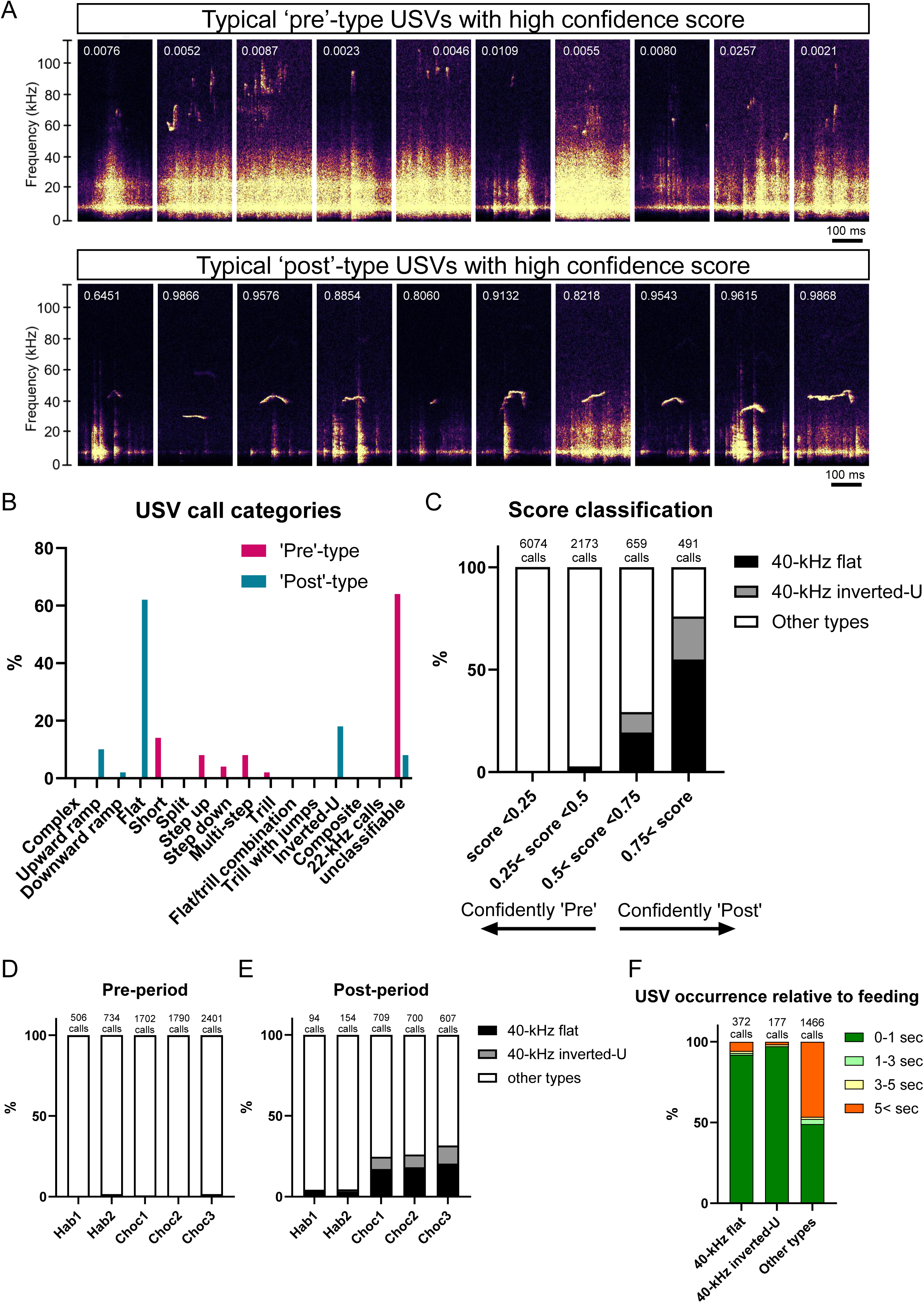
Identification of “40-kHz flat” and “40-kHz inverted-U” types as USVs emitted during chocolate consumption. A: One of the USV spectrograms with the five highest confidence scores for ‘pre’- and ‘post’-types from each of 10 rat pairs. The values shown in the figure represent logistic regression scores. Scores ranging from 0 to 0.5 are classified as ‘pre,’ while those from 0.5 to 1 are classified as ‘post.’ Additionally, values closer to 0 indicate a higher confidence in the classification as ‘pre,’ while values closer to 1 indicate a higher confidence in the classification as ‘post.’ B: Frequency of the visual classes USVs that were identified as ‘pre’- and ‘post’-type USVs by our logistic regression model. C: Percentages of 40-kHz (30-50 kHz) flat and inverted-U types among high-confidence ‘pre’ (score < 0.25), low-confidence ‘pre’ (0.25 < score < 0.5), low-confidence ‘post’ (0.5 < score < 0.75), and high-confidence ‘post’-type USVs (0.75 < score). D, E: Percentages of 40-kHz flat and inverted-U types among the total number of USVs from 10 rat pairs in the pre-(D) and post-(E) periods of each session. F: Percentages of the time intervals between 40-kHz flat, 40-kHz inverted-U, or other types of USV and the nearest instance of feeding behavior before the USV in the post-period.

**Figure 5.**
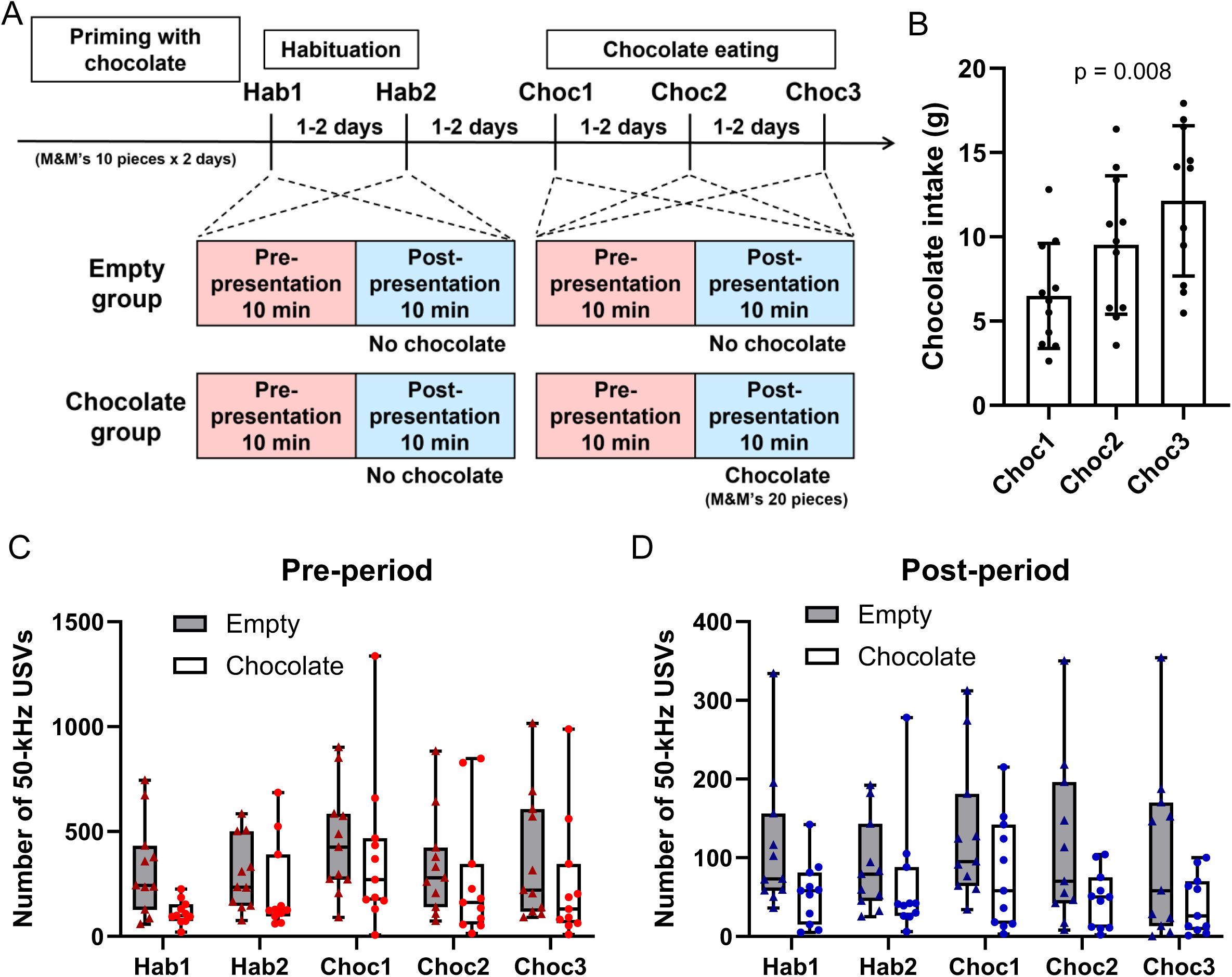
Recording of the USV test dataset. A: Schedule of USV recordings to obtain test dataset for classification using the logistic regression model. The empty group served as a control and did not receive any chocolate during the pre- or post-periods. The chocolate priming phase was conducted 2–3 days before the hab1 session, during which rats were provided with 10 pieces (approximately 10 g) of M&M’s chocolate once per day in their home cages on two occasions, with an interval of 1–2 days. Intervals between recording sessions were 1–2 days. B: Amount of chocolate intake by the chocolate group during the post-period. p = 0.008, one-way ANOVA. C, D: Number of 50-kHz short USVs during the pre-(C) and post-(D) periods.

**Figure 6.**
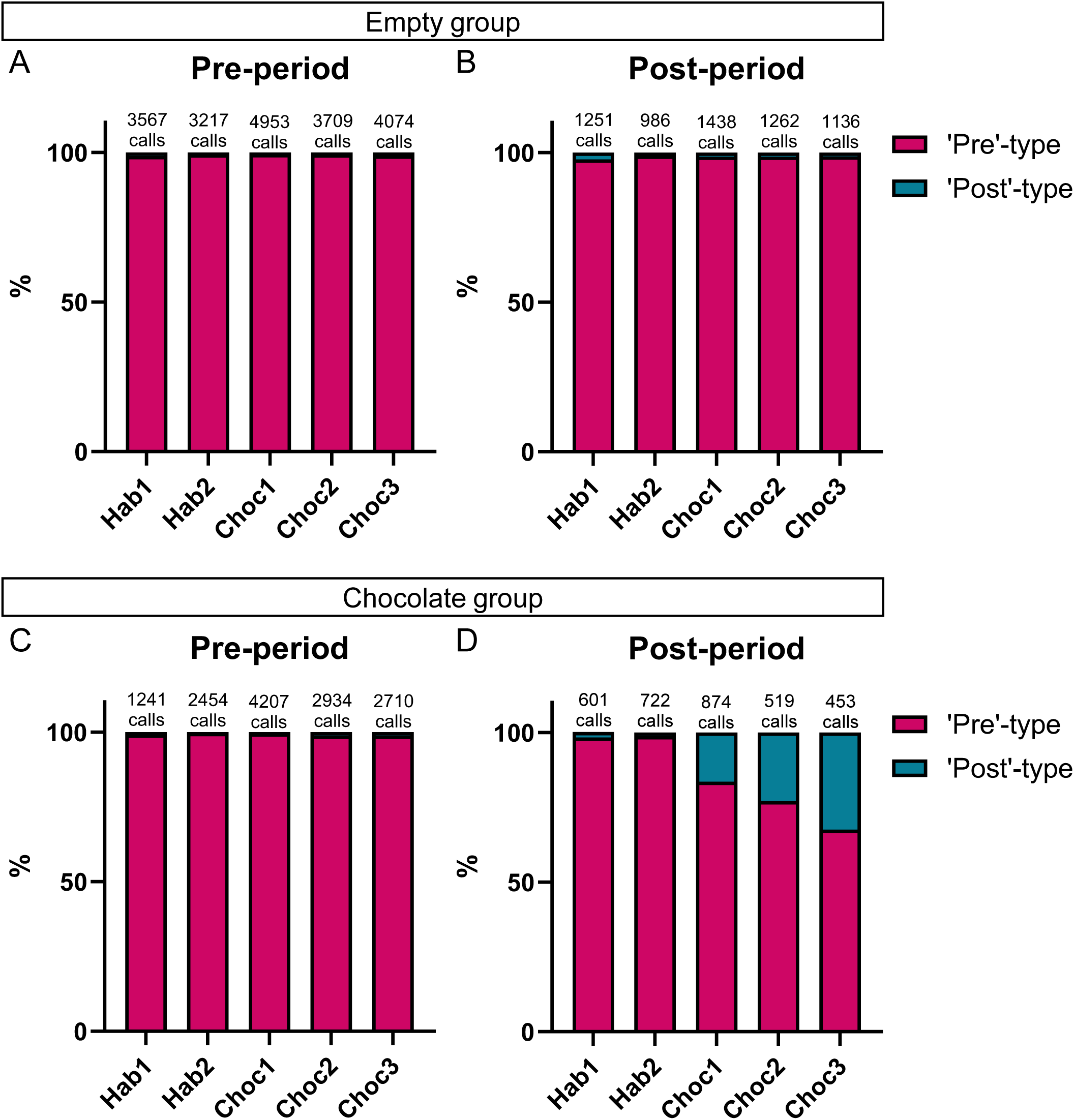
Classification of USV test dataset into ‘pre’- and ‘post’-chocolate types using the logistic regression model. A-D: Percentages of ‘pre’- and ‘post’-types among the total number of USVs from 11 rat pairs in each session. (A) Empty group in the pre-period, (B) Empty group in the post-period, (C) Chocolate group in the pre-period, (D) Chocolate group in the post-period.

**Figure 7.**
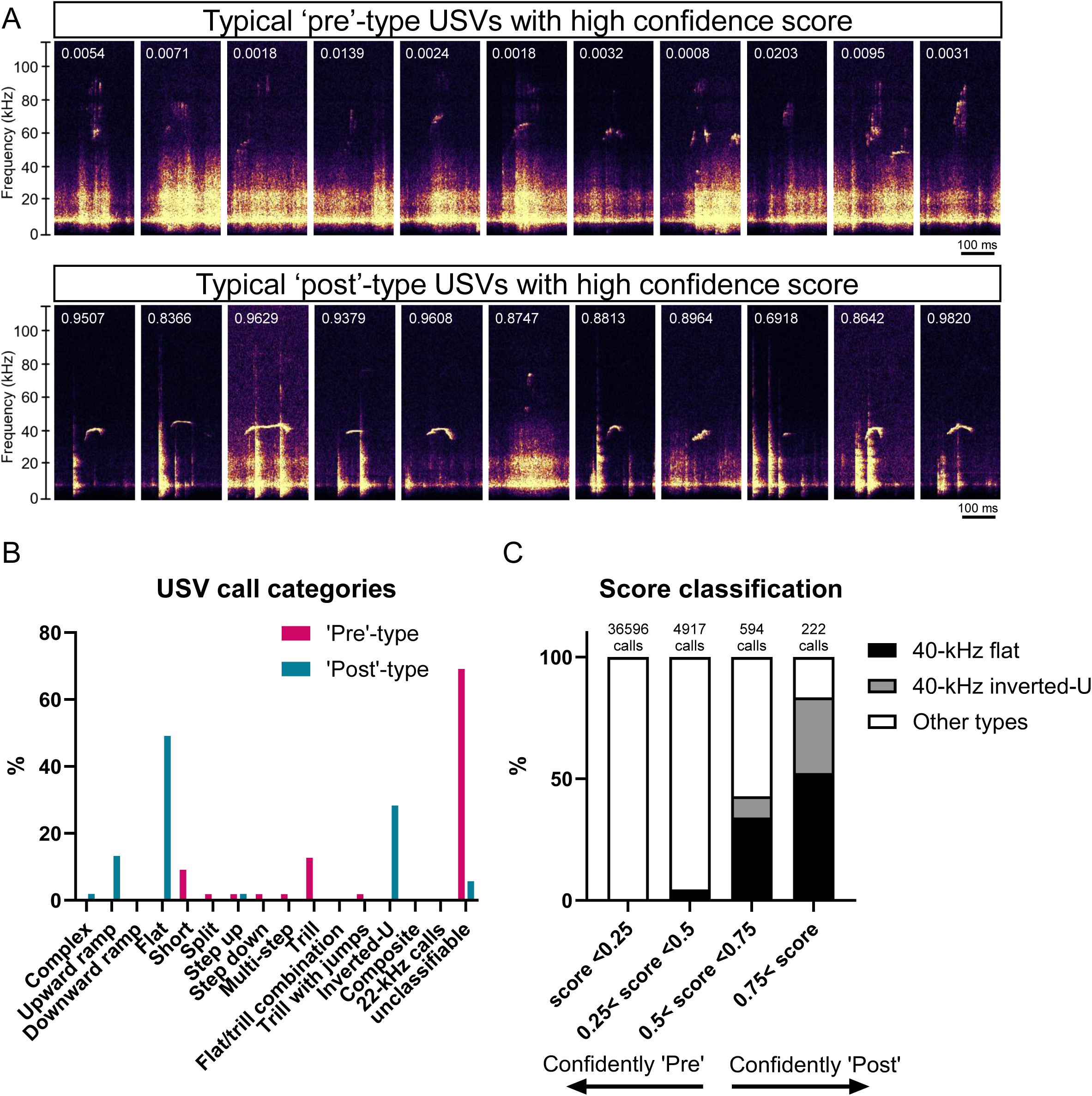
Classification of 40-kHz flat and 40-kHz inverted-U types of USVs as ‘post’-type in the test dataset. A: One of the USV spectrograms with the five highest confidence scores for ‘pre’- and ‘post’-types from each rat pair in the chocolate group. The values shown in the figure represent logistic regression scores. Scores ranging from 0 to 0.5 are classified as ‘pre,’ while those from 0.5 to 1 are classified as ‘post.’ Additionally, values closer to 0 indicate a higher confidence in the classification as ‘pre,’ while values closer to 1 indicate a higher confidence in the classification as ‘post.’ B: Frequency of visual classes of USVs identified as ‘pre’- and ‘post’-type USVs of the chocolate group. C: Percentages of 40-kHz flat (black) and inverted-U (gray) types among high-confidence ‘pre’, low-confidence ‘pre’, low-confidence ‘post’, and high-confidence ‘post’-type USVs from the entire USV test dataset.

**Figure 8.**
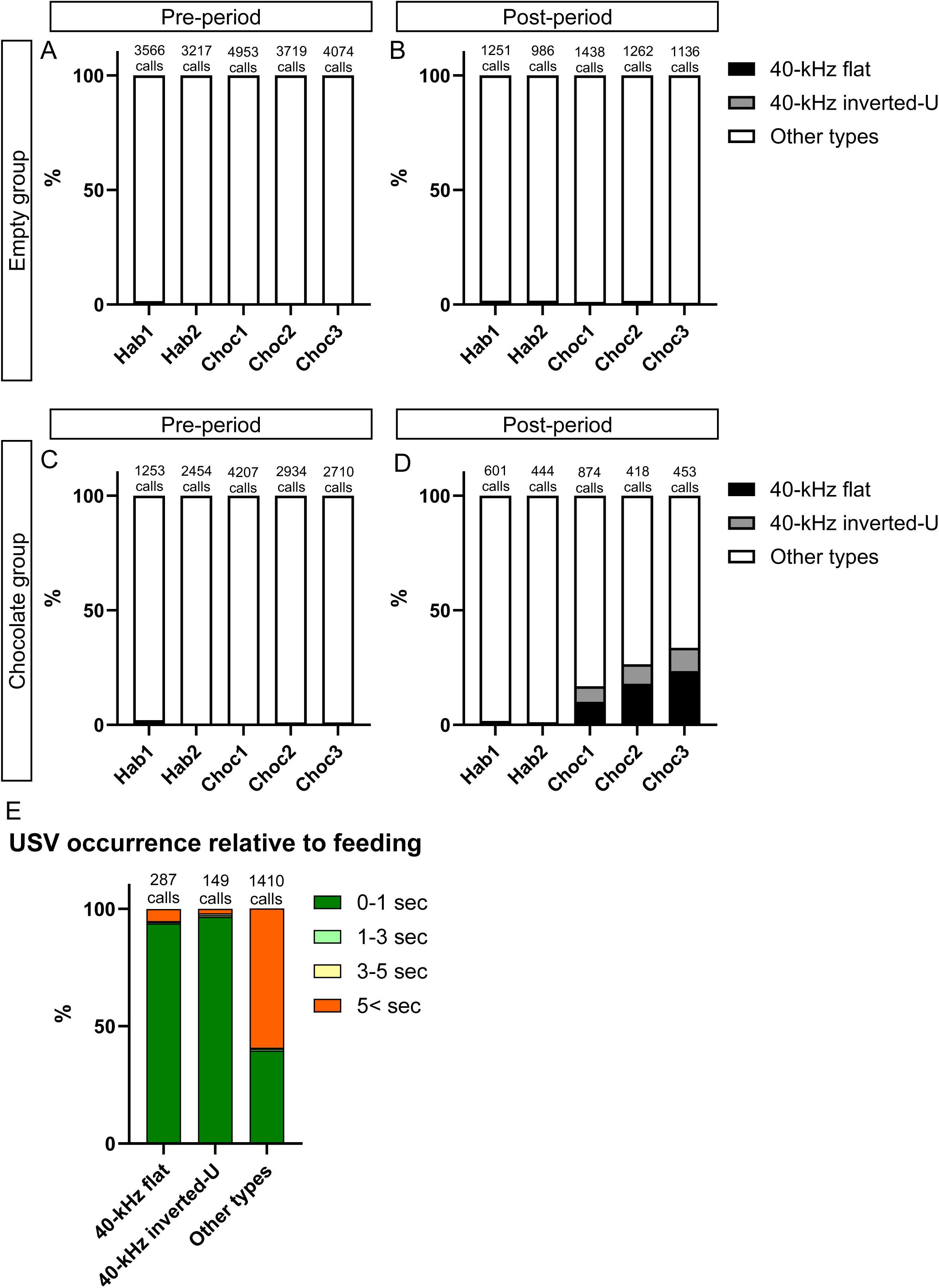
Emission of 40-kHz flat and inverted-U types of USVs during chocolate consumption in the test dataset. A-D: Percentages of 40-kHz flat and inverted-U types of USVs among all USVs from 11 rat pairs in each session. (A) Empty group in the pre-period, (B) Empty group in the post-period, (C) Chocolate group in the pre-period, (D) Chocolate group in the post-period. E: Percentages of the time intervals between 40-kHz flat, 40-kHz inverted-U, or other types of USV and the nearest instance of feeding behavior before the USV in the post-period.

**Figure 9.**
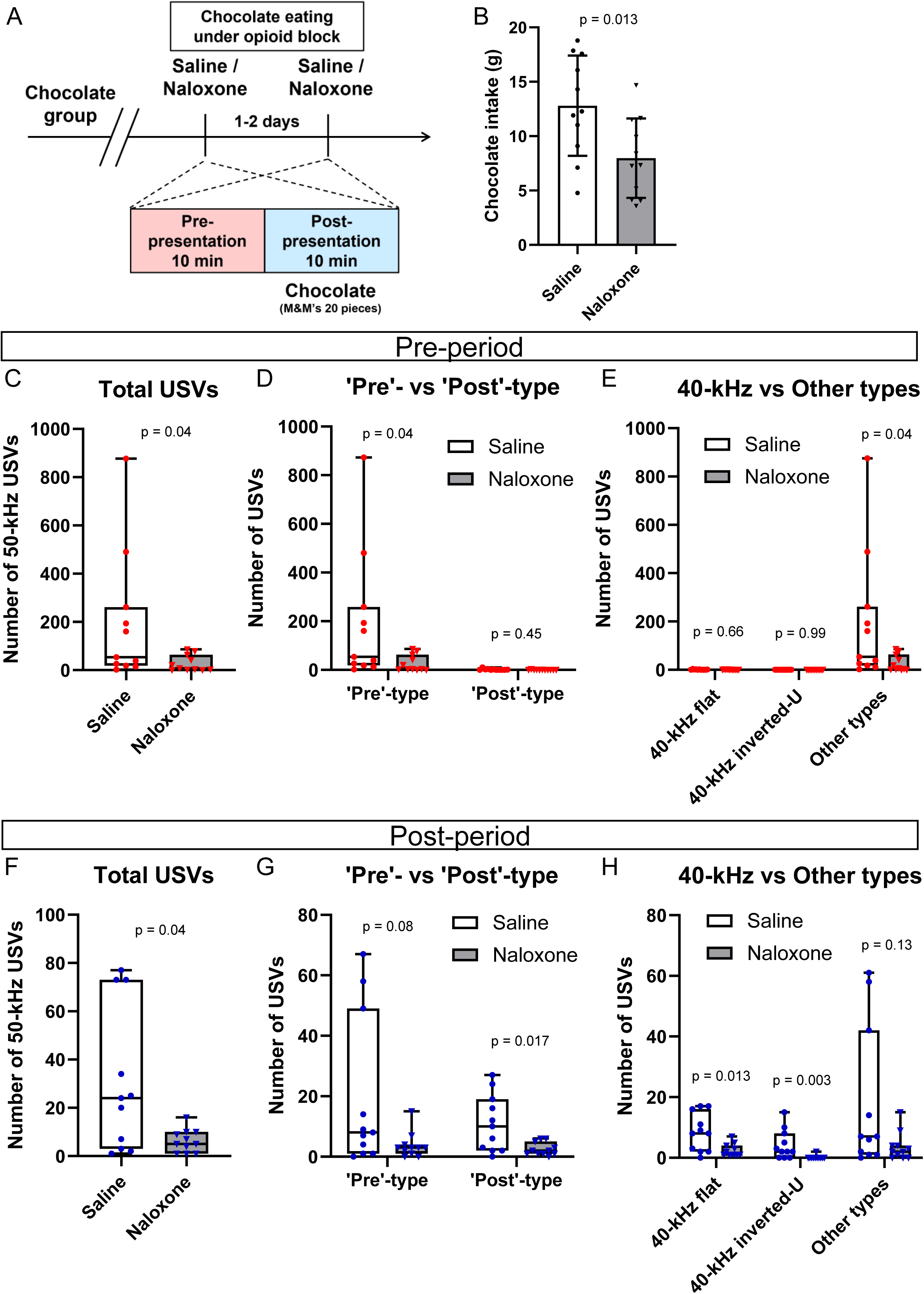
Effect of naloxone on chocolate intake and USV emission. A: Schedule of USV recording after naloxone (5 mg/kg) or vehicle saline administration. Naloxone or saline was subcutaneously injected 10 minutes before the start of USV recording. The chocolate group used for the test dataset was further subjected to this pharmacological experiment. Each rat pair received two different injections: naloxone and saline. The order of the injections was switched among the pairs for the purpose of counterbalancing. B: Amount of chocolate intake. p = 0.0013, Welch’s t test. C-H: Number of 50-kHz short USVs. (C-E) Pre-period, (F-H) post-period. (C, F) Total USVs; (D, G) ‘pre’- and ‘post’-type classification; (E, H) Number of 40-kHz flat, 40-kHz inverted-U, and USVs belonging to other type. p values were obtained from Mann– Whiteney U tests.

To optimize USV detection, experiments were conducted using pairs of rats, as previous studies have demonstrated that USV emission is more frequent in social contexts than when animals are housed individually (Inagaki et al., 2013). At the same time, the group size was limited to dyads to enable future identification of individual callers while preserving the social context. In addition, only male pairs were used to avoid confounding effects of courtship or mating behavior and to eliminate variability related to the estrous cycle, thereby ensuring a more controlled assessment of feeding-associated vocalizations.

### USV and video recording apparatus

USVs were recorded using a USB ultrasound microphone (M500-384, frequency range: 10–160 kHz; ADC resolution: 16 bits, Pettersson Elektronik AB) suspended from the ceiling of a soundproof box measuring 58 × 52 × 46 cm. There were two USB cameras (C920n, Logicool Japan, and BSW305MBK, Buffalo Inc.) positioned to capture side and back views of the rat cage, along with a red LED light to provide illumination. USV recordings were managed using BatSound Touch Lite software (Pettersson Elektronik AB), whereas OBS Studio (OBS Project) was used to simultaneously record the USV spectrograms (displayed by BatSound) and record video of the behaviors of the experimental animals from the two cameras. This setup allowed for synchronized visual inspection of both USVs and behavioral states, with minimal time discrepancy between the two.

### Recording schedule of training dataset

Fig. 1A shows the recording schedule for the training dataset, which consisted of three parts: habituation sessions, where USVs were recorded in the soundproof box without chocolate presentation (two sessions, hab1 and hab2); chocolate priming, during which rats were provided with four pieces of M&M’s milk chocolate (approximately 4 g) per day in their home cages for 3 consecutive days with no USV recording; and chocolate presentation sessions (3 sessions, choc1, choc2, and choc3). Chocolate priming was implemented to reduce neophobia towards chocolate.

Twenty minutes before the start of the recording, the bedding of the rat cages was replaced, pellet food was removed from the cage lid, and the rats were left in their home environment. Then, a pair of rats was transferred to the soundproof box in their home cage. The USVs and behaviors exhibited by the rats were recorded for the first 10 min (pre-chocolate presentation period). Then, the experimenter opened the soundproof box door and the cage lid and placed a hand into the cage (for habituation sessions) or a glass jar containing M&M’s milk chocolate (20 pieces, approximately 20 g) into the cage. The lid and door were then closed, and further recording of USVs and behaviors took place for an additional 10 min (post-chocolate presentation period). After each recording session, the rats were returned to their home environment and provided with pellet food.

### Recording schedule of test dataset

As shown in Fig. 5A, we initially conducted chocolate priming before starting USV recording, aiming to increase USV emissions during both the habituation (hab1 and hab2) and chocolate-presentation (choc1, choc2, and choc3) sessions. During priming, rats were provided with 10 pieces (approximately 10 g) of M&M’s milk chocolate per day in their home cages on two occasions with an interval of 1–2 days. We then proceeded to perform two habituation sessions, recording USV calls in the soundproof box without chocolate presentation (hab1 and hab2), followed by three chocolate presentation sessions (choc1, choc2, and choc3). The recording procedure was the same as for the training dataset, except that the empty group received an empty glass jar during the post-chocolate presentation period in the choc1, choc2, and choc3 sessions. After the choc3 session, the chocolate group underwent pharmacological experiments using naloxone (Fig. 9).

### Pharmacological experiments

To investigate the involvement of the opioid system in USV emission, we used naloxone, a nonselective competitive opioid receptor antagonist with a higher affinity for μ-opioid receptors than for κ- and δ-opioid receptors. Naloxone hydrochloride (144-09411, Wako) was dissolved in a saline solution at a concentration of 5 mg/mL. The rats received a subcutaneous injection of either naloxone or saline at a volume of 1 mL/kg body weight, administered 10 min prior to the start of USV recordings. The injection order (naloxone or saline) was alternated between pairs of rats, with six pairs receiving naloxone first, followed by saline, and five pairs receiving saline first, followed by naloxone. The recording procedure was the same as that used for the chocolate presentation sessions.

### USV detection by DeepSqueak

We used DeepSqueak v3.0.1 with MATLAB 2022b to detect 50-kHz short USVs (Coffey et al., 2019). The Rat Detector YOLO R1.mat model was utilized for USV detection, with a cutoff range of 18–100 kHz, and a rejection threshold set below 0.61. Following the automated detection and rejection of USVs, we manually eliminated pseudo-USV sounds caused by environmental noise and water intake. We also visually detected 22-kHz long USVs and excluded them from the dataset. DeepSqueak provides 11 parameters for the identified USVs, including call length, principal frequency, low frequency, high frequency, delta frequency, standard deviation of frequency, slope, sinuosity, mean power, tonality, and peak frequency. These parameters were exported to CSV files for the training of a logistic regression model and further classification of USVs.

### Supervised machine learning classification of USVs using logistic regression

We created a logistic regression model to classify USVs using the Classification Learner application in MATLAB 2022b. For the training dataset, all USVs detected by DeepSqueak during the three sessions in which chocolate was presented to 10 pairs of rats (choc1, choc2, and choc3) were used (Figs. 1 and 2). USVs emitted before the chocolate presentation (pre-period) were labeled as “pre,” whereas those emitted after the chocolate presentation (post-period) were labeled as “post.” A total of 7,909 calls were used for training, including 5,893 calls labeled as “pre” and 2,016 calls labeled as “post.” The model classified the USVs into ‘pre’-type and ‘post’-type based on 11 generated parameters (Fig. 3) and was subsequently applied to classify both the training and test datasets.

### Visual classification of USV subtypes

As shown in Figs. 4B and 7B, USVs classified as ‘pre’-type or ‘post’-type by the logistic regression model were visually reclassified by the experimenter based on conventional USV type classification using spectrogram features into 14 categories: Complex, Upward ramp, Downward ramp, Flat, Short, Split, Step up, Step down, Multi-step, Trill, Flat/trill, Trill with jumps, Inverted-U, and Composite (Wright et al., 2010). USVs that did not fall into any of these categories were labeled as “unclassifiable.” For each pair of rats in the training dataset (10 pairs) and the chocolate group in the test dataset (11 pairs), the top five USVs with the highest confidence scores for being either ‘pre’-type or ‘post’-type were chosen for visual classification.

We then visually examined all USVs from both the training and test datasets and classified them into three types: 40-kHz flat type (near-constant frequency with a mean slope between −0.2 and 0.2 kHz/ms, with a principal frequency range of 30–50 kHz), 40-kHz inverted-U type (a monotonic increase followed by a monotonic frequency decrease, each of at least 5 kHz, with a principal frequency range of 30–50 kHz), or other type (Figs. 4C–E, 7C, 8A–E, and 9E, H).

### Visual classification of feeding behavior

Feeding behaviors associated with USV occurrence during the post-period of chocolate presentation sessions (choc1, choc2, and choc3) were assessed visually by the experimenter using the USB camera recordings. For each USV, the time interval (in seconds) from the most recent instance in which either or both rats exhibited chocolate biting or mastication behavior to the occurrence of that USV was measured. The intervals were then classified into four categories: 0–1 sec, 1–3 sec, 3–5 sec, and 5< sec (Figs. 4F and 8E).

### Statistical analyses

For datasets subjected to statistical analysis, normality was initially assessed using the Shapiro–Wilk test (JASP 0.8.1.0). When all datasets followed a normal distribution, data were presented as means with standard deviations using bar charts, and parametric tests were applied. If any dataset deviated from normality, data were presented as medians with interquartile ranges using box-and-whisker plots, and non-parametric tests were used. All statistical analyses were conducted using GraphPad Prism 10. The number of replicates was determined by referencing similar previous studies and is specified for each experiment in the Results section. The outcomes of the statistical tests are provided in Table S1.

### Data availability statement

The data that support the findings of this study, including USV audio files (WAV format), behavioral video files (MP4 format), USV detection files generated by DeepSqueak (MAT format), and statistical analysis files, are available from the corresponding author, Koshi Murata, upon reasonable request. The logistic regression model will be shared via an online repository after acceptance.

## Results

### Increased emission of 50-kHz short USVs during both pre- and post-chocolate presentation periods

To investigate whether rats emit specific subtypes of USVs when consuming palatable foods, we recorded USVs during chocolate feeding. Adult male Sprague–Dawley rats were housed in pairs and placed in a soundproof box for USV recording, where they were allowed access to M&M’s milk chocolate. Each recording session consisted of a 10-min pre-presentation period (pre-period), during which the rats did not have access to the chocolate, followed by a 10-min post-presentation period (post-period). To habituate the rats to the soundproof box and establish baseline vocalization levels, two habituation sessions (hab1 and hab2) were carried out before chocolate presentation. During these sessions, the door of the soundproof box and the lid of the cage were opened at the end of the pre-period to simulate chocolate delivery and habituate the rats to the experimenter’s hand intrusion without presenting any chocolate. After the habituation sessions, the rats were primed with four pieces of M&M’s chocolate (approximately 4 g per day for 3 consecutive days) in their home environment. Following this, three sessions of chocolate presentation (choc1, choc2, and choc3) were carried out. During these sessions, the home cages containing each rat pair were placed in the soundproof box, and their USVs were recorded for 10 min (pre-period). Subsequently, 20 pieces of chocolate (approximately 20 g) were provided to the rats, and their USVs were recorded for another 10 min (post-period). An interval of 1–2 days was maintained between the recording sessions (Fig. 1A). Rat behaviors were video-recorded, and chocolate intake was measured. Using DeepSqueak, we identified 50-kHz (30–100 kHz) short USVs from the recorded audio data. Notably, fasting was not part of the experimental protocol.

Chocolate consumption increased slightly across the three chocolate presentation sessions but not significantly (one-way ANOVA, F_(2, 27)_=1.365, *p=*0.27; Fig. 1b). During the pre-period of the two habituation sessions, most rat pairs emitted a low number of 50-kHz short USVs (Fig. 1C). In contrast, there was a significant increase in USVs during the pre-period of the three sessions that included chocolate presentation (Kruskal–Wallis test, *p=*0.0104; Fig. 1C). Similarly, during the post-period of the two habituation sessions, most rats emitted few USV calls, but a significant increase in USV calls was observed during those sessions that included chocolate presentation (Kruskal– Wallis test, *p=*0.0012, Fig. 1D).

### Distinct USV spectrograms between pre- and post-chocolate presentation periods

We then proceeded to examine whether the spectrograms of the 50-kHz short USVs emitted during the post-period when chocolate was available differed from those emitted during the pre-period (when chocolate was unavailable). We conducted a statistical analysis that compared 11 parameters (call length, principal frequency, low frequency, high frequency, delta frequency, standard deviation of frequency, slope, sinuosity, mean power, tonality, and peak frequency) of the USV spectrograms measured by DeepSqueak between the pre- and post-periods of three sessions in which chocolate was presented. There was a total of 5,893 USV calls in the pre-period and 2,016 USV calls in the post-period from the sessions choc1, choc2, and choc3. All 11 parameters exhibited significant differences between the pre- and post-periods, as determined by the Mann–Whitney *U* test (Fig. 2A–K). USVs emitted during the post-period were characterized by lower frequency, longer duration, increased loudness, and improved signal-to-noise ratio compared to those emitted during the pre-period.

To obtain representative examples of USV spectrograms from both the pre- and post-periods, we selected five USVs that were closest to the median values of the 11 parameters in each period. The USVs from the pre-period showed various shapes of spectrograms in the 50–60 kHz range. In contrast, the USVs from the post-period exhibited inverted-U, flat, and upward ramp shapes in the 40–50 kHz range (Fig. 2L).

To further investigate if the differences in the 11 parameters were consistent among individual rat pairs within sessions, we conducted separate Mann–Whitney U tests for each pair. The results are presented in Fig. 2M, organized from high to low chocolate consumption across the three chocolate presentation sessions (from left to right). Fewer parameters exhibited statistically significant differences between the pre- and post-periods among individual pairs during the habituation sessions (hab1 and hab2). However, during the chocolate presentation sessions (choc1, choc2, and choc3), rat pairs with high chocolate consumption exhibited marked differences in vocalization between the pre- and post-sessions.

### Classification of ‘post’-type USVs using a supervised machine learning model, highlighting 40-kHz flat and 40-kHz inverted-U types of USVs during chocolate-feeding

To identify USV subtypes unique to the post-period, we then examined if a supervised machine learning model could distinguish between USVs as ‘pre’-type or ‘post’-type by analyzing differences in the parameters of USV spectrograms between the pre- and post-periods. A logistic regression model was developed using the 11 parameters presented in Fig. 2A–K to categorize USVs as either ‘pre’-type or ‘post’-type. The training dataset included USVs from three chocolate presentation sessions, comprising all calls in the pre-period (n = 5,893 calls) labeled as ‘pre-period type’ and all calls in the post-period (n = 2,016 calls) as ‘post-period type’. The resulting model achieved an accuracy of 82.5% and an area under the curve (AUC) of 0.78 (Fig. 3A). The model primarily emphasized call length and tonality (Fig. 3B). Another model was constructed using a dataset with randomized pre-period and post-period labels, which yielded an accuracy of 83.1% and an AUC of 0.51 (Fig. 3A), classifying all USVs as ‘pre’-type. The higher AUC in the original model suggests that USVs in the post-period have distinct spectrograms compared to those in the pre-period and that our model effectively used these differences to classify USVs into the ‘pre’- and ‘post’-types.

We applied the classifier model to the USVs emitted during both the two habituation sessions and the three chocolate presentation sessions, which were utilized as training datasets. The number of USVs classified as ‘pre’- and ‘post’-types and the ratio of both types among the total USVs were assessed. During the pre-period of both the habituation and chocolate presentation sessions, most USVs were classified as ‘pre’-type (Fig. 3C). In contrast, ‘post’-type USVs made up a larger percentage (40–46%) of the USVs emitted during the chocolate presentation sessions (Fig. 3D). These results suggest that ‘post’-type USVs were seldom emitted in the absence of chocolate but became more prevalent when chocolate was available to the rats.

Next, we analyzed the spectrogram of USVs classified as ‘pre’-type and ‘post’-type. From each pair of rats, we selected the top five USVs with the highest classification confidence, as shown in Fig. 4A and Figure S1, and evaluated them visually based on conventional USV-type categories (Wright et al., 2010). The ‘pre’-type USVs with high confidence scores were mainly short or unclassifiable. Conversely, most of the ‘post’-type USVs with high confidence scores belonged to the flat or inverted-U types, accounting for 62% and 18%, respectively (Fig. 4B). Besides the spectrogram patterns, a distinct feature of ‘post’-type USVs was their frequency range (Fig. 2B). Both flat and inverted-U types exhibited a principal frequency range within 40 kHz (30–50 kHz; Fig. 4A, lower case). This 40-kHz frequency range has been reported in previous studies conducted during the consumption of a standard laboratory diet (Takahashi et al., 2010; Champeil-Potokar et al., 2023).

Next, we evaluated the proportion of 40-kHz flat and 40-kHz inverted-U type USVs within those classified as ‘pre’- and ‘post’-type by the model (Fig. 4C). The ‘pre’-type USVs contained few 40-kHz flat or 40-kHz inverted-U types. In contrast, among the USVs classified as ‘post’-type with high confidence (logistic regression score: 0.75<), 55% were 40-kHz flat USVs and 21% were 40-kHz inverted-U USVs. Among the USVs classified as ‘post’-type with low confidence (logistic regression score: between 0.5-0.75), 19% were 40-kHz flat USVs and 10% were 40-kHz inverted-U USVs. Since the ‘post’-type USVs occurred when chocolate was presented to the rats, we hypothesized that 40-kHz flat and 40-kHz inverted-U USVs were emitted when chocolate was available. We visually assessed all USVs to determine whether they were 40-kHz flat or 40-kHz inverted-U types (Fig. 4D and E). Both 40-kHz flat and 40-kHz inverted-U USVs occurred when chocolate was available, specifically during the post-periods of the chocolate presentation sessions (choc1, choc2, and choc3).

We further investigated whether the 40-kHz flat and inverted-U USVs were associated with actual chocolate consumption. Using video recordings, we analyzed the behaviors of the rats and measured the time intervals between each USV and the nearest instance of feeding behavior, defined as biting or mastication by either one or both rats before the USV (Fig. 4F). In summary, 91.9% of 40-kHz flat USVs and 97.2% of 40-kHz inverted-U USVs were emitted within 0-1 second of a feeding behavior, whereas 49.0% of other USVs occurred within this time window. In contrast, 46.5% of other USVs were emitted more than 5 seconds after a feeding event. These results suggest that 40-kHz flat and inverted-U USVs are tightly coupled with chocolate-feeding behavior.

### Applicability of the logistic regression model and reproducibility of the 40-kHz flat and inverted-U USVs during chocolate consumption

We next validated the logistic regression model to distinguish between the ‘pre’- and ‘post’-type USVs in different rat pairs. We also investigated the association between the ‘post’-type, 40-kHz flat, and 40-kHz inverted-U USVs and chocolate consumption. We created 11 new rat pairs and included another group in which empty containers (without chocolate) were presented in the post-period (empty group, 11 pairs; Fig. 5A). Furthermore, we included chocolate priming before the habituation sessions and increased the amount of chocolate used for priming (10 g × 2 days) for both chocolate and empty groups. The chocolate consumption of the chocolate group significantly increased across the three sessions (one-way ANOVA, F_(2, 30)_=5.66, *p=*0.008; Fig. 5B). Both the empty and chocolate groups exhibited a larger number of 50 kHz USVs during the two habituation sessions and the three chocolate presentation sessions (Fig. 5C and D) than the previous dataset (Fig. 1C and D). We speculate that the increased number of USV calls reflects a heightened expectation level for the chocolate of rats, resulting from the additional amount provided during priming.

Next, we categorized all USVs detected in this test dataset as ‘pre’- and ‘post’-types using the classification model (Fig. 3A and B). During the pre-period, nearly all USVs produced in both the habituation and chocolate presentation sessions of both the empty and chocolate groups were classified as ‘pre’-types (97.7–99.7%, Fig. 6A–C). In contrast, there were significant occurrences of the ‘post’-type USVs only during the post-period of the three sessions in the chocolate group (specifically when chocolate was accessible to the rats; Fig. 6D).

We then analyzed the spectrogram of USVs identified as ‘pre’-type and ‘post’-type in this new dataset. From each pair of rats in the chocolate group, we selected the top five USVs with the highest classification confidence (Fig. 7A and Figure S2) and evaluated them visually based on the conventional USV-type classification (Wright et al., 2010). The ‘pre’-type USVs with high confidence scores were mostly short or unclassifiable types, whereas most of the ‘post’-type USVs with high confidence scores were either flat or inverted-U types, representing 49% and 28%, respectively (Fig. 7B). Both the flat and inverted-U types had a principal frequency range within 40 kHz (30–50 kHz).

Next, we examined the proportion of 40-kHz flat and 40-kHz inverted-U type USVs among those classified as ‘pre’- and ‘post’-type by the model in the new dataset. The ‘pre’-type USVs contained few 40-kHz flat or 40-kHz inverted-U types. In contrast, among the USVs classified as ‘post’-type with high confidence (logistic regression score: 0.75<), 52% were 40-kHz flat USVs, and 31% were 40-kHz inverted-U USVs. For USVs classified as ‘post’-type with low confidence (logistic regression score: between 0.5-0.75), 34% were 40-kHz flat USVs, and 9% were 40-kHz inverted-U USVs (Fig. 7C). We then visually inspected all USVs in the new dataset to determine whether they belonged to the 40-kHz flat or 40-kHz inverted-U type. The results revealed that both 40-kHz flat and 40-kHz inverted-U USVs occurred when chocolate was available, specifically during the post-period of the chocolate presentation sessions in the chocolate group (Fig. 8A–D).

We then examined whether the 40-kHz flat and inverted-U USVs were associated with chocolate consumption in the new dataset, as depicted in Fig. 4F. Again, 93.7% of 40-kHz flat USVs and 96.6% of 40-kHz inverted-U USVs were emitted within 0-1 second of a feeding behavior, whereas 39.6% of other USVs occurred within this time window. In contrast, 59.3% of other USVs were emitted more than 5 seconds after a feeding event. These results further confirmed the close association between 40-kHz flat and inverted-U USVs and chocolate-feeding behavior.

### Effect of opioid blocker naloxone on 40-kHz flat and inverted-U USVs during chocolate consumption

We next examined the influence of opioids on ‘post’-type, 40-kHz flat, and 40-kHz inverted-U USVs, given that opioid receptor blockers reduce ‘liking’ reactions in the taste reactivity test (Parker et al., 1992). Following three chocolate presentation sessions with the chocolate group, the naloxone injection experiment was conducted with the same 11 rat pairs (Fig. 9A). The rats were administered either naloxone (5 mg/kg) or vehicle saline subcutaneously, and after 10 min, USV recordings and chocolate presentation were carried out in the same manner as in the previous sessions. Naloxone significantly decreased chocolate consumption (Welch’s t-test, t_(19.01)_=2.72, *p=*0.013; Fig. 9B).

Overall, USV emissions were reduced by naloxone during both the pre- and post-periods (Mann–Whitney *U* test, *p=*0.04 for pre-period, *p=*0.04 for post-period; Fig. 9C and F). When examining the effect of naloxone on ‘pre’- and ‘post’-type USV emissions, ‘pre’-type USVs showed a significant decrease in the pre-period (*p=*0.04), whereas ‘post’-type USVs did not show a significant change (*p=*0.45; Fig. 9D). In the post-period, ‘pre’-type USVs were not significantly reduced by naloxone (*p=*0.08), but ‘post’-type USVs were significantly decreased (*p=*0.017; Fig. 9G). Furthermore, we analyzed the effect of naloxone on 40-kHz flat and inverted-U USV emissions. In the pre-period, naloxone did not lead to a significant decrease in 40-kHz flat and inverted-U USVs (*p=*0.66 for flat USVs, *p=*0.99 for inverted-U USVs) but decreased other types of USVs (*p=*0.04; Fig. 9E). In contrast, in the post-period, naloxone significantly reduced both 40-kHz flat and inverted-U USVs (*p=*0.013 for flat USVs, *p=*0.003 for inverted-U USVs) but did not significantly decrease other types of USVs (*p=*0.13; Fig. 9H). These results suggest that overall, 50-kHz short USV emissions are linked to the opioid system and that 40-kHz inverted-U and flat USVs during chocolate consumption are particularly responsive to naloxone.

## Discussion

In this study, we investigated whether specific subtypes of USVs are emitted by rats when they consume palatable food, such as chocolate, and whether these USVs can be objectively classified using machine learning. Through analysis of acoustic parameters of USVs and the application of a logistic regression model, we found that rats emit distinct USV subtypes when given access to chocolate, particularly 40-kHz inverted-U subtypes, in addition to 40-kHz flat subtypes which are known to occur during the consumption of standard food pellets. These findings raise the possibility that rat USVs are modulated by food palatability. Moreover, the emission of both 40-kHz inverted-U and flat USVs was significantly reduced following administration of the opioid receptor antagonist naloxone, suggesting that the opioid-regulated reward system contributes to their emission.

### Modulation of USVs by food palatability

Our findings regarding the emission of 40-kHz ranged USVs during chocolate consumption are partially consistent with previous reports (Takahashi et al., 2010; Champeil-Potokar et al., 2023). According to Takahashi et al., USVs in Sprague–Dawley rats can be classified into three main clusters based on frequency range and call duration (Takahashi et al., 2010). USVs emitted during the consumption of standard laboratory chow generally fall within the 40-kHz range (30–50 kHz). The spectrogram shape of these USVs is predominantly flat, with the inverted-U type being rarely observed (accounting for less than 1% relative to flat USVs). In our study, the occurrence of inverted-U USVs during chocolate consumption was markedly higher, representing approximately 48% (training dataset) and 45% (test dataset) of the number of flat USVs. Champeil-Potokar et al. reported that Wistar rats emit USVs while masticating food (Champeil-Potokar et al., 2023). In their study, using standard laboratory chow, the predominant USV type was flat in the 40-kHz range (35–50 kHz), with very few inverted-U types observed. In our spectrogram analysis, we observed similar overlap between sounds produced during chocolate mastication and USVs, identifying both flat and inverted-U USVs within the 40-kHz range (30–50 kHz). These results suggest that rats typically produce flat USVs in the 40-kHz range while feeding, and that palatability of the food may modulate their vocalizations from flat to inverted-U shape in the spectrogram.

Evidence from food-associated calls in rhesus macaques also supports the idea that food palatability influences vocal expression (Hauser and Marler, 1993a, 1993b). Rhesus macaques exhibit different types of vocalizations when encountering rare, highly preferred foods, such as coconut, compared to standard chow. Further research is required to clarify this phenomenon by feeding rats different types of food with varying degrees of palatability (tasteless, low-nutrient, high-fat, or high-sugar) and comparing the resulting USV responses.

### Machine learning-based classification of USV subtypes

A methodological advancement in our study is the use of machine learning to classify USV subtypes emitted before and after chocolate presentation periods based on their spectrogram characteristics. We utilized a logistic regression model for supervised binary classification, which facilitated confidence scoring that helped identify USV subtypes related to chocolate consumption. Conventional methods for USV classification involve manual visual inspection, which is time-consuming, possibly biased, and not easily scaled for large datasets. In contrast, our logistic regression model, trained on 11 spectrogram parameters, achieved high-throughput classification with an accuracy of 82.5% and an AUC of 0.78. This approach facilitated a more objective and consistent analysis across different rat pairs and experimental sessions, reducing human error and inter-observer variability.

To evaluate the effectiveness of our model, we created another logistic regression model using a randomized training dataset with shuffled pre- and post-labeling and compared the AUC values. This approach yielded a model with a very low AUC value (0.51), indicating poor classification performance with the randomized dataset. Although the model with the randomized dataset achieved a seemingly higher accuracy (83.1%) than our original model (82.5%), this higher accuracy was likely due to the imbalance in the ratio of pre- and post-labeled USVs in the training dataset (pre-label: 5,893 calls; post-label: 2,016 calls). In fact, the model with the randomized dataset classified all USVs in the training dataset as ‘pre’-type. These results suggest that USVs in the post-period have distinct spectrograms compared to those in the pre-period and that our model successfully utilized these differences to classify USVs into the ‘pre’- and ‘post’-types.

Our model identified call length and tonality as the most important features distinguishing USVs emitted before and after chocolate presentation. During the post-period, USVs were generally longer than those in the pre-period. Furthermore, rats remained still while feeding chocolate, which reduced noise from the bedding being rubbed and resulted in higher tonality, specifically in the signal-to-noise ratio. Our model effectively captured these USV characteristics and related behavioral states, ensuring the reliability of machine learning for USV classification.

### Roles of opioids in USV emissions during palatable feeding and their implications for positive emotional states

In this study, we found that systemic administration of naloxone significantly reduced the emission of both 40-kHz flat and 40-kHz inverted-U USVs during chocolate consumption, while having less impact on other USV types. This suggests that the emission of these two subtypes is modulated by the endogenous opioid system. The opioid system is well known to contribute to positive emotional states in animals (Emery and Akil, 2020; Murata et al., 2024). In particular, during the hedonic “liking” reactions observed in the taste reactivity test, opioid receptor antagonists reduce characteristic responses such as rhythmic tongue protrusions (Parker et al., 1992).

Previous studies have also shown that administration of psychostimulants and opioid blockers modulates the emission of 50-kHz short USVs and their subtypes, implicating contributions of opioids to vocalization patterns linked to positive motivational states (Burgdorf et al., 2001; Buck et al., 2014). Our findings extend this framework by demonstrating that palatable food–associated 40-kHz inverted-U and flat USVs are likewise sensitive to opioid receptor blockade, indicating that these vocalizations may serve as behavioral markers of positive emotional states during feeding. Future research should aim to identify the specific neural circuits responsible for the generation of these opioid-sensitive USVs.

### Limitations and future directions

Our study was designed to minimize the influence of social and sexual factors, focusing specifically on a simplified and controlled feeding context. To this end, we used male rat dyads that had been pair-housed for weeks, allowing the establishment of a stable social relationship prior to experimentation. While this design allowed us to isolate chocolate-feeding–related USV patterns under controlled conditions, it also raises the question of how generalizable these findings are to more naturalistic or socially complex environments. Future studies should explore whether similar USV patterns, particularly the 40-kHz inverted-U subtype, are observed under more ecologically valid conditions, including group housing, individually housed (solo) rats, or female subjects.

Another important limitation concerns the identification of the vocalizing individual. Although our behavioral analysis demonstrated that the inverted-U and flat USVs were more tightly linked to chocolate-feeding behavior than other USVs, we could not distinguish which of the two rats in a pair produced each vocalization. As such, the current data cannot rigorously determine whether the emitting rat was directly engaged in feeding at the moment of USV emission. To establish whether the 40-kHz inverted-U USV serves as a real-time emotional marker of feeding pleasure, future work must incorporate improved tracking of individual vocalizers and concurrent behavioral states.

The identification of a novel 40-kHz inverted-U USV subtype associated with chocolate consumption adds to our understanding of the biological significance of USV diversity. As shown in our data, these USVs were tightly linked to chocolate-feeding behavior and were suppressed by opioid receptor blockade, suggesting their potential involvement in affective processes related to palatable food intake. USVs are widely recognized as signals for social communication, and in this context, it is worth considering whether the 40-kHz inverted-U subtype identified here may also play a communicative role (Wöhr, 2017). Hauser and Marler reported that rhesus monkeys produced distinct vocalizations when discovering highly palatable foods, which functioned as honest signals that reduced aggression from conspecifics and increased food access (Hauser and Marler, 1993a, 1993b). In rodents, social transmission of food preferences has been demonstrated: observer rats acquire a preference for foods consumed by demonstrators, primarily through olfactory cues (Galef and Whiskin, 1992; Bessières et al., 2017). These findings raise the intriguing possibility that food-related USVs—such as the 40-kHz inverted-U subtype—may not only reflect internal affective states but also broadcast information about the presence or value of palatable food to nearby conspecifics. Future studies are needed to test this hypothesis by examining the emission and reception of such USVs under socially dynamic conditions.

In summary, our findings suggest that specific USV subtypes, particularly the 40-kHz inverted-U type, may serve as acoustic signatures during palatable food consumption. These vocalizations are modulated by the opioid system and can be efficiently detected using machine learning. Extending this framework to other sensory modalities or types of reward will be essential for evaluating the generalizability and neurobiological basis of vocal-affective expressions in animals.

## Conflict of interest statement

The authors declare no competing interests.

## Acknowledgements

We thank Hideki Yoshikawa, Eri Murai, Takako Maegawa, Mayumi Yamamoto, and the members of the Fukazawa Laboratory and the Life Science Research Laboratory at the University of Fukui for their technical and administrative assistance.

---

**Supplementary Figure 1.**
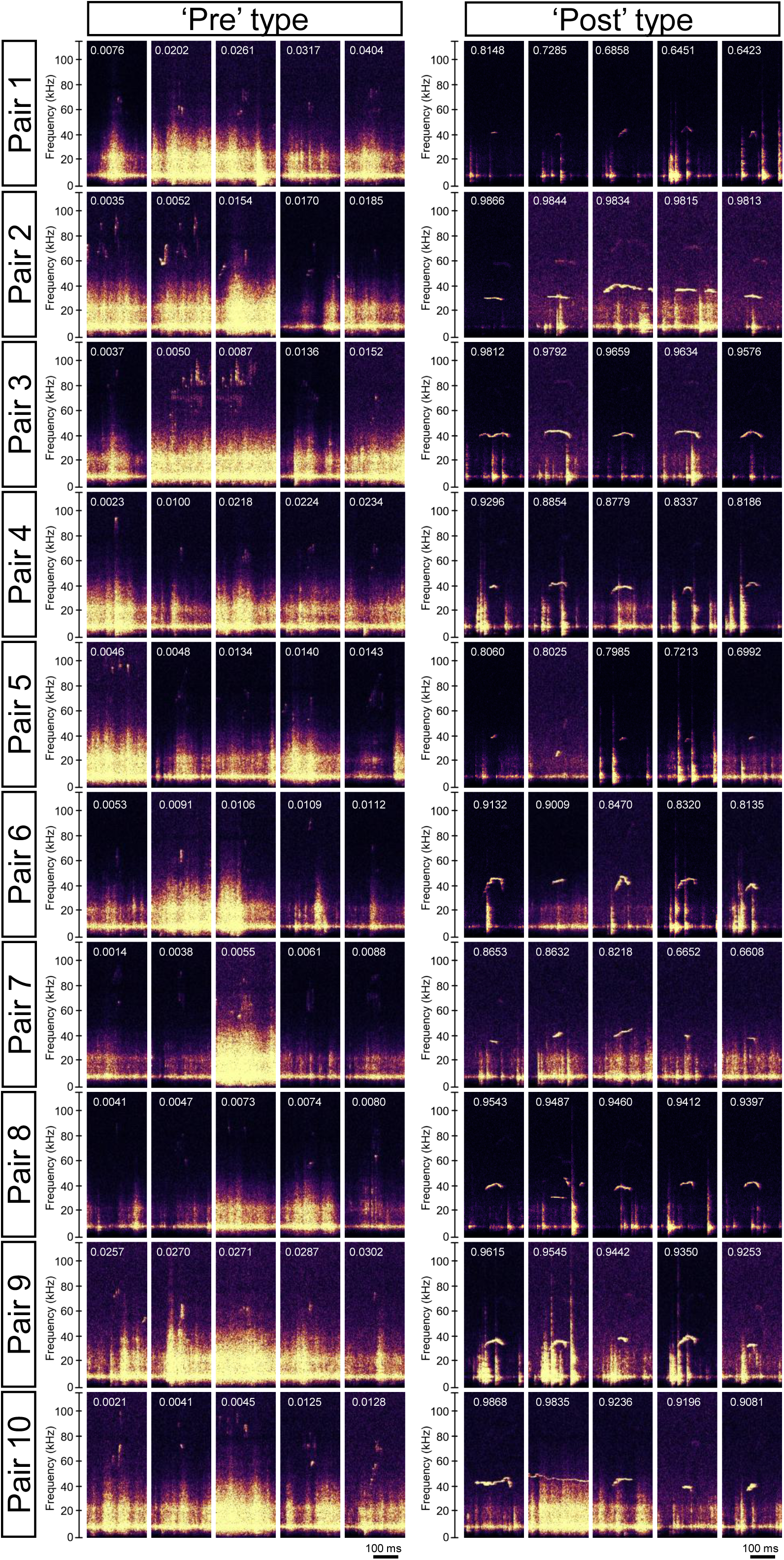
Representative ‘pre’- and ‘post’-type USVs of the training dataset. Five USVs with the highest confidence score from each of the 10 pair for classification as ‘pre’- or ‘post’-type were selected. The values shown in the figure represent logistic regression scores. Scores ranging from 0 to 0.5 are classified as ‘pre,’ while those from 0.5 to 1 are classified as ‘post.’ Additionally, values closer to 0 indicate a higher confidence in the classification as ‘pre,’ while values closer to 1 indicate a higher confidence in the classification as ‘post.’

**Supplementary Figure 2.**
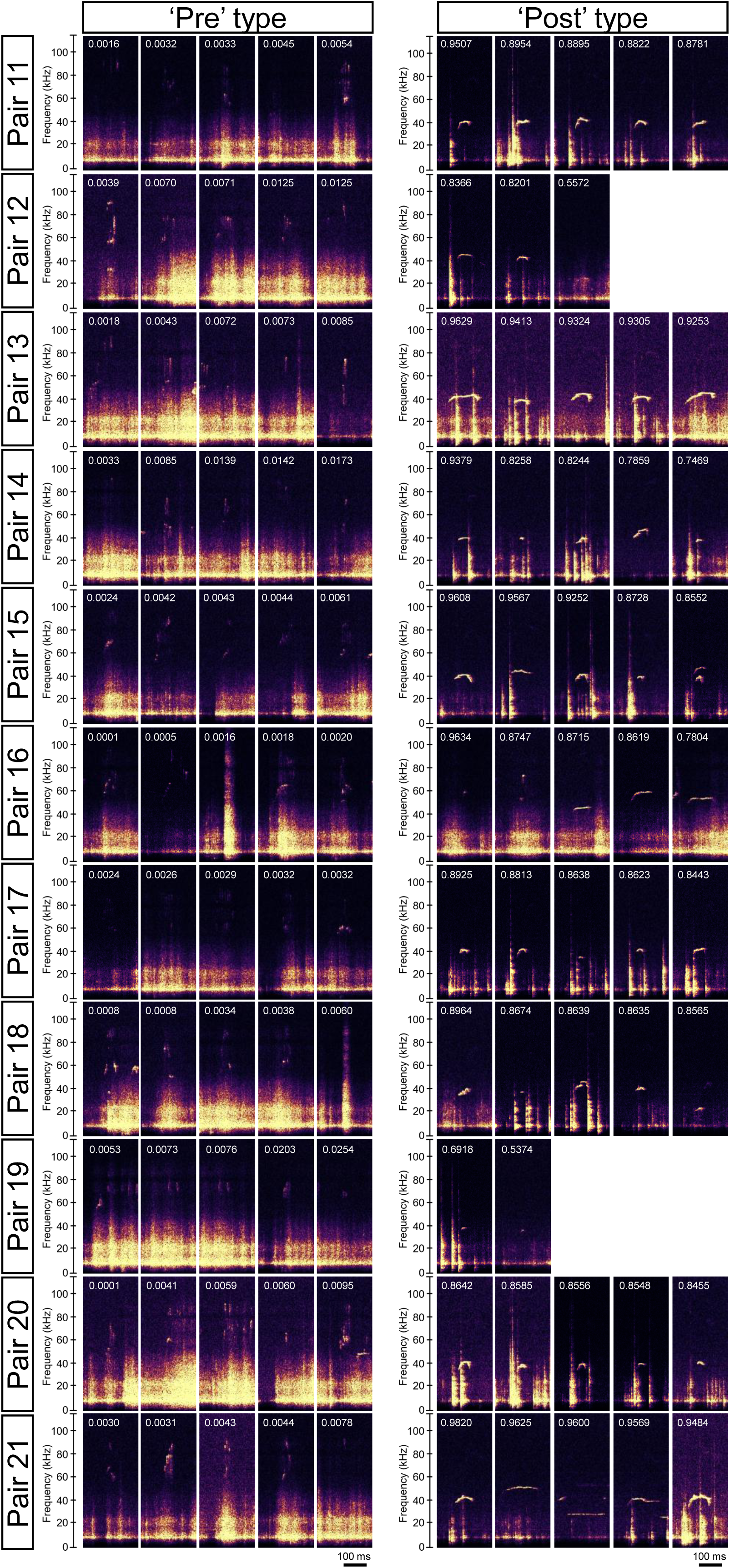
Representative ‘pre’- and ‘post’-type USVs of the test dataset. Five USVs with the highest confidence score from each of the 11 pair in the chocolate group for classification as ‘pre’- or ‘post’-type were selected. For Pairs 12 and 19, fewer than five ‘post’-type USVs were available. The values shown in the figure represent logistic regression scores. Scores ranging from 0 to 0.5 are classified as ‘pre,’ while those from 0.5 to 1 are classified as ‘post.’ Additionally, values closer to 0 indicate a higher confidence in the classification as ‘pre,’ while values closer to 1 indicate a higher confidence in the classification as ‘post.’

**Supplementary Table 1.** Summary of statistical tests.

| Figure | Description | Analysis | t, F, U, H value | p value | Sample size |
| --- | --- | --- | --- | --- | --- |
| 1B | chocolate intake | one-way ANOVA | F(2, 27) = 1.365 | p = 0.27 | n = 10 pairs per group |
| 1C | number of USVs in pre-period | Kruskal-Wallis | H = 13.18 | p = 0.0104 | n = 10 pairs per group |
| 1D | number of USVs in post-period | Kruskal-Wallis | H = 18.01 | p = 0.0012 | n = 10 pairs per group |
| 2A | Call length | Mann-Whitney | U = $4.548 \times 10^6$ | p < 0.001 | n = 5893 calls (pre), 2016 calls (post). |
| 2B | Principal frequency | Mann-Whitney | U = $8.655 \times 10^6$ | p < 0.001 | n = 5893 calls (pre), 2016 calls (post). |
| 2C | Low frequency | Mann-Whitney | U = $8.446 \times 10^6$ | p < 0.001 | n = 5893 calls (pre), 2016 calls (post). |
| 2D | High frequency | Mann-Whitney | U = $9.004 \times 10^6$ | p < 0.001 | n = 5893 calls (pre), 2016 calls (post). |
| 2E | Delta frequency | Mann-Whitney | U = $6.797 \times 10^6$ | p < 0.001 | n = 5893 calls (pre), 2016 calls (post). |
| 2F | Frequency standard deviation | Mann-Whitney | U = $7.030 \times 10^6$ | p < 0.001 | n = 5893 calls (pre), 2016 calls (post). |
| 2G | Slope | Mann-Whitney | U = $6.832 \times 10^6$ | p < 0.001 | n = 5893 calls (pre), 2016 calls (post). |
| 2H | Sinuosity | Mann-Whitney | U = $6.562 \times 10^6$ | p < 0.001 | n = 5893 calls (pre), 2016 calls (post). |
| 2I | Mean power | Mann-Whitney | U = $3.955 \times 10^6$ | p < 0.001 | n = 5893 calls (pre), 2016 calls (post). |
| 2J | Tonality | Mann-Whitney | U = $3.342 \times 10^6$ | p < 0.001 | n = 5893 calls (pre), 2016 calls (post). |
| 2K | Peak frequency | Mann-Whitney | U = $8.709 \times 10^6$ | p < 0.001 | n = 5893 calls (pre), 2016 calls (post). |
| 5B | chocolate intake | one-way ANOVA | F(2, 30) = 5.657 | p = 0.0082 | n = 11 pairs per group |
| 9B | chocolate intake | Welch's t test | t(19.01) = 2.724 | p = 0.014 | n = 11 pairs per group |
| 9C | Total USVs in pre-period | Mann-Whitney | U = 29.5 | p = 0.042 | n = 11 pairs per group |
| 9D | 'pre'-type USVs in pre-period | Mann-Whitney | U = 29.5 | p = 0.042 | n = 11 pairs per group |
|  | 'post'-type USVs in pre-period | Mann-Whitney | U = 50.0 | p = 0.45 | n = 11 pairs per group |
| 9E | 40-kHz flat USVs in pre-period | Mann-Whitney | U = 53.0 | p = 0.66 | n = 11 pairs per group |
|  | '40-kHz inverted-U USVs in pre-period | Mann-Whitney | U = 60.5 | p = 0.99 | n = 11 pairs per group |
|  | other type USVs in pre-period | Mann-Whitney | U = 29.0 | p = 0.039 | n = 11 pairs per group |
| 9F | Total USVs in post-period | Mann-Whitney | U = 30.0 | p = 0.044 | n = 11 pairs per group |
| 9G | 'pre'-type USVs in post-period | Mann-Whitney | U = 34.0 | p = 0.084 | n = 11 pairs per group |
|  | 'post'-type USVs in post-period | Mann-Whitney | U = 25.0 | p = 0.017 | n = 11 pairs per group |
| 9H | 40-kHz flat USVs in post-period | Mann-Whitney | U = 23.5 | p = 0.013 | n = 11 pairs per group |
|  | '40-kHz inverted-U USVs in post-period | Mann-Whitney | U = 19.5 | p = 0.0035 | n = 11 pairs per group |
|  | other type USVs in post-period | Mann-Whitney | U = 37.5 | p = 0.13 | n = 11 pairs per group |

## Notes

### Competing Interest Statement

The authors have declared no competing interest.

### Funding

- Lotte Research Foundation
- Urakami Foundation for Food and Food Culture Promotion
- Takeda Science Foundation
- Japan Society for the Promotion of Science: 17KK0190, 20H05955, 21H05817, 21K06440, 24K02131

